# AstraPTM2: A Context-Aware Transformer for Broad-Spectrum PTM Prediction

**DOI:** 10.1101/2025.10.03.680341

**Authors:** Çağlar Bozkurt, Alexandra Vasilyeva, Aniruddh Goteti

## Abstract

Post-translational modifications (PTMs) are covalent changes in proteins after biosynthesis that shape stability, localization, and function. While numerous computational tools exist for PTM site prediction, most struggle to handle full-length proteins without truncating context, focus on only a limited number of PTM types, and perform unevenly on rare modifications.

We present AstraPTM2, a transformer-based model that predicts 39 distinct PTM types on full-length sequences. By combining ESM-2 embeddings, AlphaFold2-derived structural features, and protein-level descriptors, AstraPTM2 captures both short-range motifs and long-range dependencies. Training uses a three-stage curriculum and adaptive focal loss to balance rare and common PTMs, followed by per-label affine calibration and optimized thresholds for well-calibrated probabilities.

In hold-out tests, AstraPTM2 achieves AUROC = 0.99 and macro-F1 = 59% across 39 PTM types, with particularly strong performance on rare motif-driven PTMs such as O-linked glycosylation and sumoylation. Results are available through the Orbion web platform, which offers synchronized 2D and 3D visualizations, dual prediction modes (calibrated and exploratory), and reproducible exports to support downstream experimental planning.

AstraPTM2 can be accessed at https://www.orbion.life.

## Introduction

Post-translational modifications (PTMs) are covalent changes occurring after protein synthesis. Common examples include phosphorylation (adding a phosphate group), glycosylation (adding sugar molecules) and ubiquitination (usually tagging for degradation). PTMs are key to a protein’s stability, localization and interactions, as well as metabolic regulation throughout the cell cycle (1). For this reason, PTM sites can be targeted for rational mutant design; for example, glycosylation sites are often removed for better protein crystallization (2). Research has also highlighted the clinical significance of PTM prediction, particularly in cancer and neurodegenerative diseases, where abnormal PTM profiles can reveal therapeutic targets. By identifying key phosphorylation or ubiquitination sites, drug design can focus on inhibiting or modifying these PTMs, thereby improving treatment specificity and efficacy.

By altering how proteins fold and function, PTMs modulate nearly every aspect of cellular biology and can play pivotal roles in disease progression. However, PTMs can be difficult to detect experimentally (3). This is further complicated by the fact that PTM pathways vary by organism and cell type, and some modifications are exclusive to eukaryotes, which poses challenges to selecting suitable expression systems. In addition, many PTMs are reversible and can only be detected under specific conditions or in certain cell cycle stages. Consequently, reliable *in silico* PTM prediction is an important tool for protein research.

Numerous computational tools have been developed to predict PTM sites from protein sequence and structural cues. Some approaches focus on proximity-based features, while others draw on evolutionary history to highlight residues likely to undergo modification. Early computational tools leveraged structural data (e.g., solvent accessibility) and family-level modification counts to predict PTM sites; subsequent models incorporated PTM co-occurrence patterns and protein structural information. Several models have been recently published that evaluate multiple PTM labels, such as MIND-S (4), PTM-Mamba (5), PDeepPP (6), DeepMVP (3), MTPrompt-PTM (9), MeToken (10), PTM-CMGMS (11), PTM-GPT2 (12), SiteTACK (13).

Despite the notable progress in the field of PTM prediction, considerable challenges remain. One key issue is the reliance on limited context windows, which restricts the ability to capture long-range dependencies. Additionally, many models do not include a sufficiently large or diverse set of proteins in their training corpora and only address a small number of best-studied PTMs. While important for some applications, these approaches are not applicable for assessing a broad range of PTMs.

To address these gaps, we present AstraPTM2, a context-aware transformer model that:

- Predicts 39 PTM types on full-length proteins — the broadest coverage to date, without truncating context windows.
- Balances rare and common PTMs through curriculum learning and per-label calibration within a single unified model.
- Captures both local motifs and long-range dependencies via multi-scale convolutions and attention.
- Is deployed in a production-ready web platform featuring synchronized 2D and 3D views and reproducible, exportable results.

## Methods

This section describes the dataset construction, model architecture, training regime, and calibration procedure for AstraPTM2. We summarize the input features, outline the backbone and auxiliary tasks, and detail the curriculum learning strategy that improved generalization across 39 PTM labels.

### Training Data Structure

The training data were processed from publicly available research studies, enriched with additional embeddings and features. Each protein sequence was represented by a set of heterogeneous descriptors:

- **ESM-2 Embeddings**: residue-level embeddings of shape [S, 1280], generated from the pretrained esm2 t33 650M UR50D model (7).
- **AlphaFold2 Structural Block (AF2 Block)**: per-residue features of shape [S, 6], including normalized pLDDT, relative solvent accessibility (RSA), 3-state secondary structure (H/E/C) one-hot encoding, and a has-structure mask. The AF2 structures were retrieved from the AlphaFold Protein Structure Database (8).
- **Orbion Enrichments**: a combination of protein- and residue-level descriptors [1, 70], which includes but not limited to molecular weight, isoelectric point, GRAVY score, sequence length, and number of bonds. These provide additional contextual information about the protein.
- **Amino Acid Representation**: a one-hot encoding of the 20 canonical amino acids.
- **Relative Position Encoding**: an extra column was appended to each sequence, scaled from 0→1 along sequence length.

After combining these sources, the final input per protein was a residue-level feature matrix of size [S, 1377].

To ensure balanced coverage across PTMs, a subset of 100,000 proteins was created from the curated database of 237,813 proteins. Rare PTM labels were oversampled, while a greedy gain strategy selected proteins that maximized label coverage. The resulting set was shuffled and split into 70% training, 15% validation, and 15% test; which then was oversampled to 108,291 proteins, which was used only for the training set, but not on validation or test sets.

**Figure 1:**
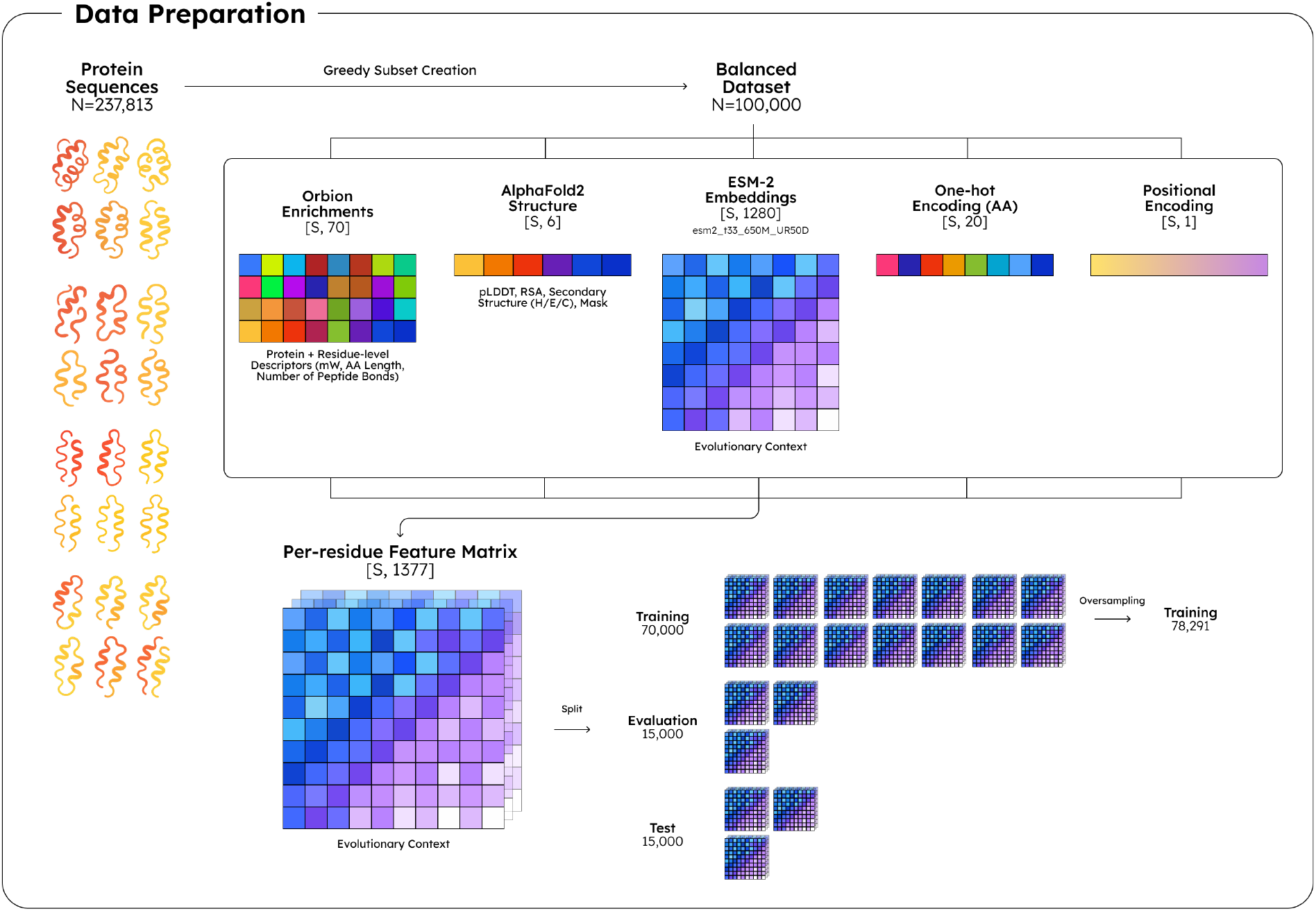
Data preparation flow for the dataset used to train, evaluate, and test the model

### Architecture

The model is a **single-backbone, multi-task, residue-level PTM classifier**. It integrates sequence, structural, and enrichment features, then applies a Transformer encoder augmented with **multi-scale convolutions** and a **global gating mechanism**.

Outputs are partitioned into **label groups** (common, medium, rare) with separate linear heads, while a dedicated **phosphorylation tri-head** predicts STY, HK, and ANY phosphorylation signals. Auxiliary tasks provide additional supervision: site-window classification, kinase-group classification, and optionally a structural graph encoder branch from AF2 contact graphs.

**Table 1:**
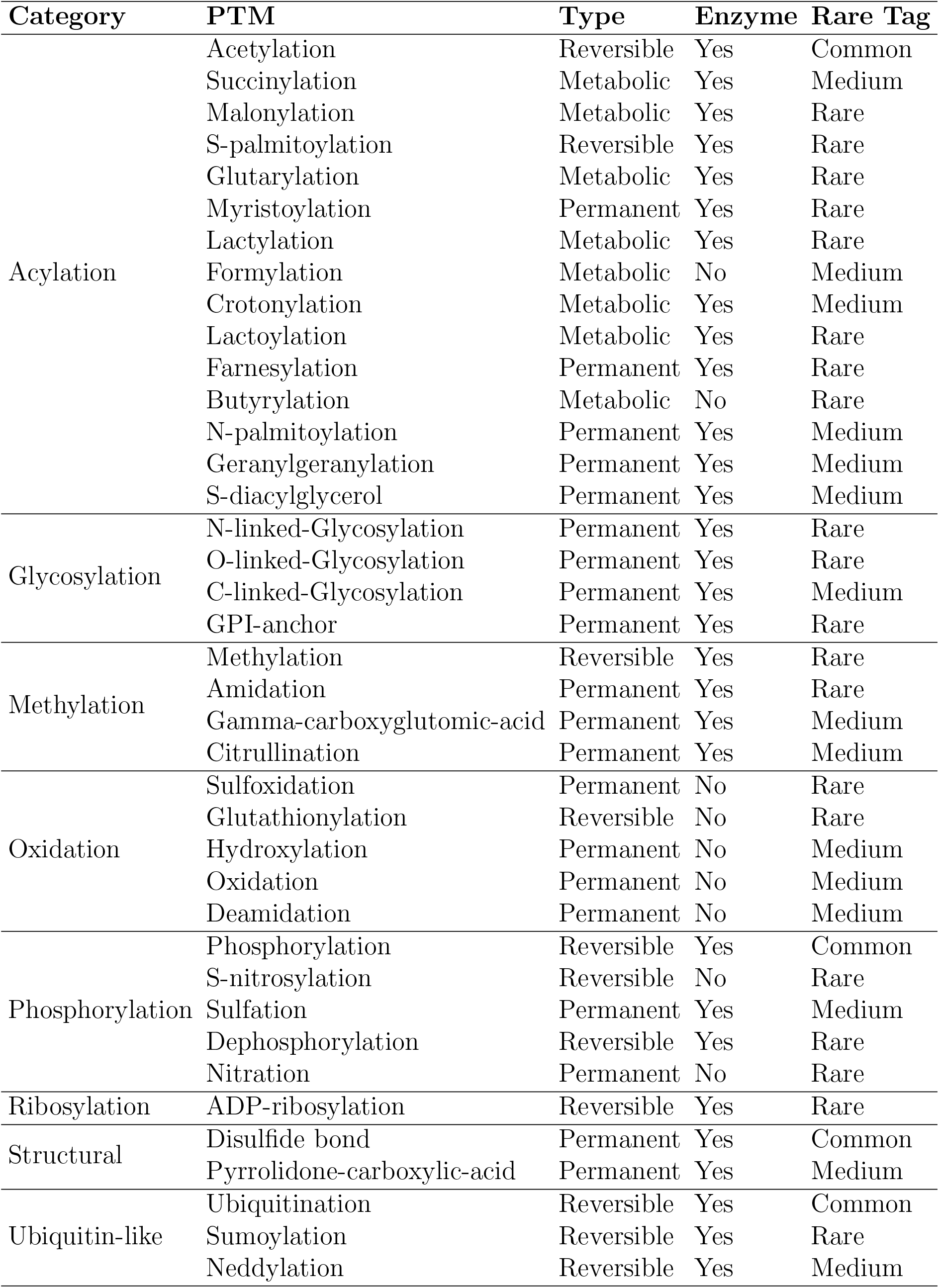
Overview of 39 PTM labels assessed in AstraPTM2.

Training is distributed with accelerate (fp16, DDP) and uses **adaptive focal loss** with per-label weighting (*α, γ*, pos weight). Class imbalance is handled through a **three-stage**

**curriculum** that progressively introduces rarer labels:

- **Stage A (Common Only):** 25 epochs, backbone unfrozen, LR(head/backbone)=4e-4/4e-4.
- **Stage B (Common + Medium):** 15 epochs, backbone frozen, LR=3e-4/3e-5.
- **Stage C (All Labels):** 15 epochs, backbone unfrozen, LR=1e-4/3e-5.

Each stage uses separate optimizers for backbone vs heads, checkpoints after completion, and supports fast resume.

### Encoder & Context

The encoder consists of:

- **LayerNorm** → Linear projection (1377→768).
- **8 Transformer Blocks**: multi-head attention with 24 heads using Rotary Positional Embeddings (RoPE), followed by GELU MLPs and residual connections.
- **Global Gating**: mean-pooled sequence vector g modulates token states through sigmoid(Wg).
- **Multi-Scale Convolutional Context**: three depth-wise convolutions (dilated k=7, k=5, k=3), summed and added back residually.

### Heads and Auxiliary Tasks

- **Grouped Heads:** one linear head per label group (common, medium, rare). Logits are rescaled by learnable temperature multipliers (T mul) and gamma multipliers (*γ* mul).
- **Phospho Tri-Head:** three outputs per residue (STY, HK, ANY back-off).
- **Auxiliary Heads:** Site-window classification: pooled subsequences → MLP → total out logits (*λ*=0.5); Kinase-group classification: global vector g → Linear → 5-way logits (4 families + other).
- **Optional Structural Branch:** a two-layer AF2 contact-graph encoder (GCN-Conv) adds structural context; when PyG is unavailable, a mean neighbor aggregation is used instead.

### Calibration and Thresholds

After training, the model outputs are calibrated to improve operating thresholds:

- **Affine Per-Label Calibration:** logistic regression fits (a, b) per label, producing probabilities p = sigmoid(a·logit + b).
- **Threshold Search:** a two-stage procedure, first coarse (step 0.02), then fine (*±* 0.03 range with step 0.005). Beta-smoothed priors are applied: *α*=*β*=1 for common labels, 0.5 otherwise.
- **Global Fallback:** if per-label *τ* cannot be determined, a single global *τ* is chosen to maximize macro-F1.

### Training Strategy

Our training strategy evolved through successive iterations aimed at balancing label distribution, architectural complexity, and curriculum scheduling. Early baselines on large datasets (230K, 75K proteins) with the new architecture produced relatively low F1 scores (*<*11%), highlighting the strong influence of class imbalance and noisy negatives.

To address this, we integrated several strategies:

- **Curriculum Learning:** Training was staged (common → medium → rare), with experiments on varying epoch allocations (e.g., 10-15-15, 25-15-15, 30-15-15). Longer emphasis on the common group provided stability, while extending rare-label phases improved generalization. The best configurations added over +10 net points on the macro-F1 metric.
- **Sampling & Windows:** Negative sampling ratios and residue window sizes were tuned. Increasing window size to 35–40 and setting NEG SAMPLING RATIO between 1.0–1.5 yielded better contextual balance, boosting F1 by up to +6 points.
- **Loss Dynamics:** Multiple loss functions were tested: focal *γ* decay, dynamic *γ*/*α* schedules, positive–unlabeled loss, class-balanced focal, and asymmetric focal. Among these, adaptive focal loss with *γ* decay consistently improved minority label learning, while asymmetric focal collapsed.
- **Architectural Enhancements:** We iteratively introduced **multi-scale depthwise convs, tri-mix context, branch-specific gating, residue-type mixers**, and structural branches (GCN, GAT). These improved local motif capture and structure awareness, with some runs surpassing **55% macro-F1**.
- **Calibration:** Post-training calibration was refined via per-label affine scaling and *β*-smoothed *τ* searches. Proper feasibility filtering and group-wise multipliers stabilized evaluation, often adding +2–3 F1, and ultimately pushed performance into the **high 50s**.

Overall, the training strategy matured into a combination of balanced manifests, curriculum scheduling, multi-scale context encoders, and per-label calibrated thresholds, supported by extensive ablations.

**Figure 2:**
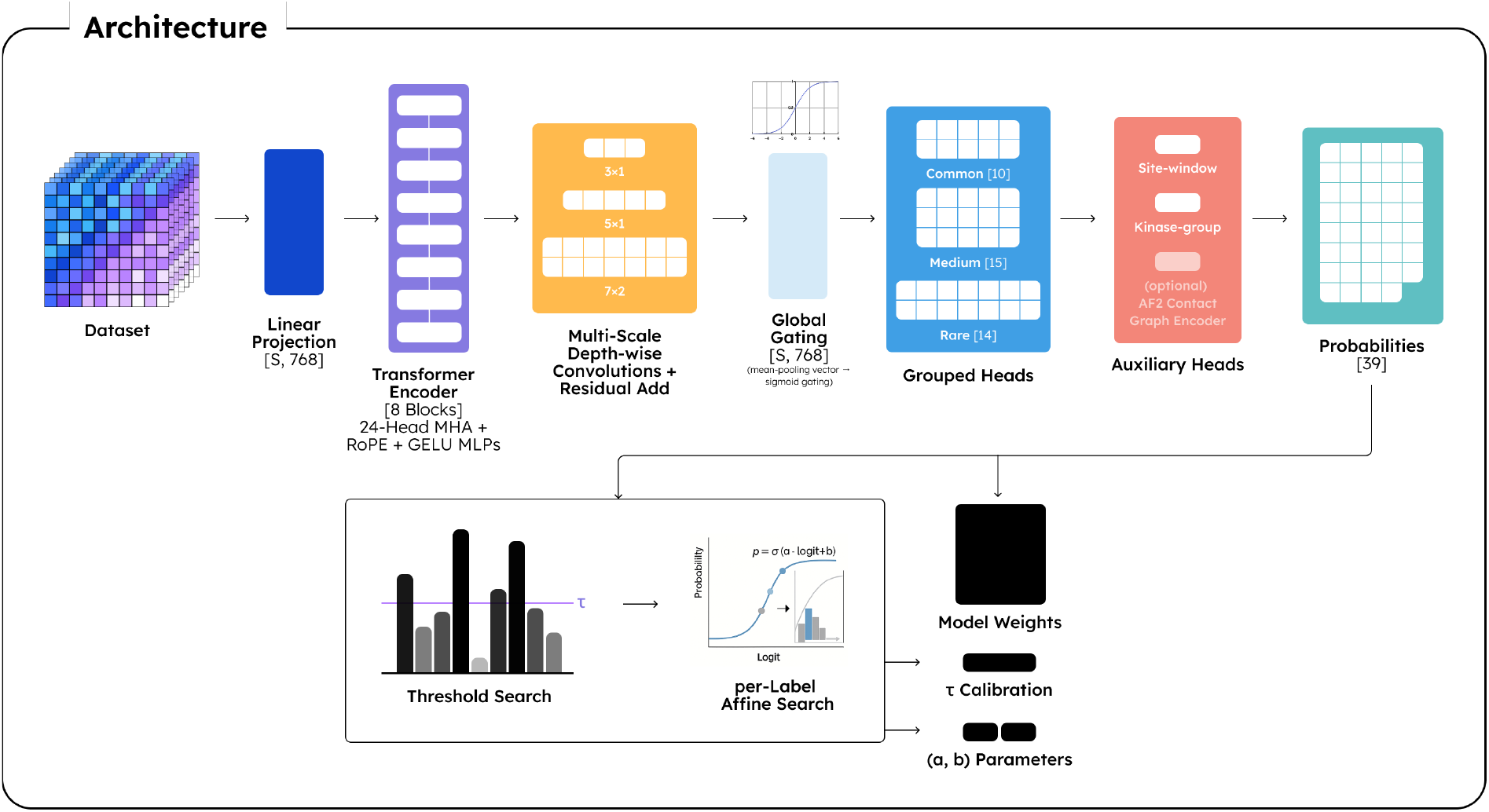
The training model architecture and the outputs

**Figure 3:**
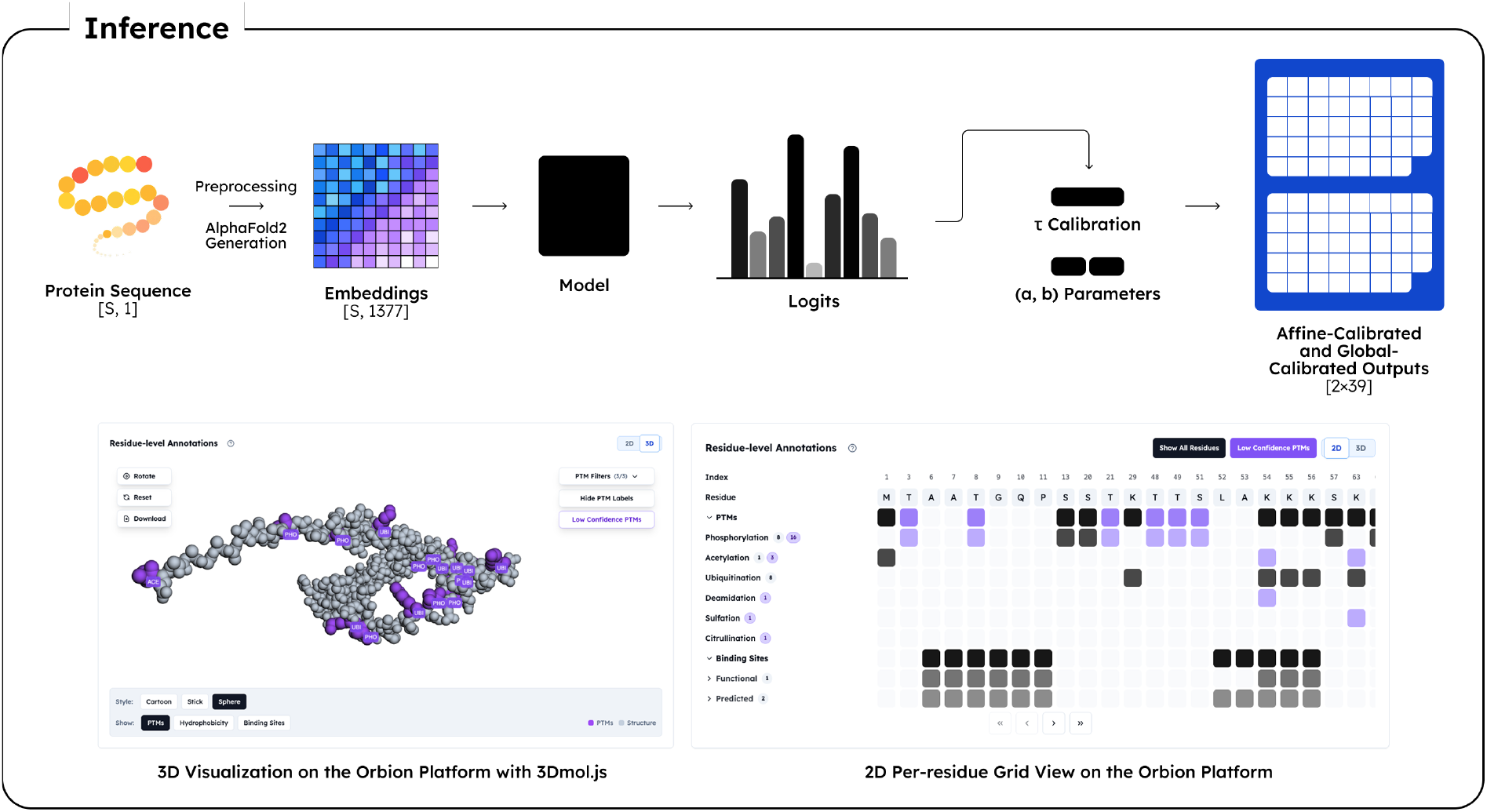
The inference flow for the PTM prediction requests

## Results

### Performance Metrics

The final model reached a macro-F1 of 59% across 39 PTM types, a major improvement over early baselines (*<*20%). AUROC was 0.99 for the entire dataset and remained consistently high for separate labels (0.95 median), indicating strong ranking ability even where thresholded classification was weaker.

When separated by PTM, most labels showed good performance (see Figure 4) and can be grouped as follows:

**Figure 4:**
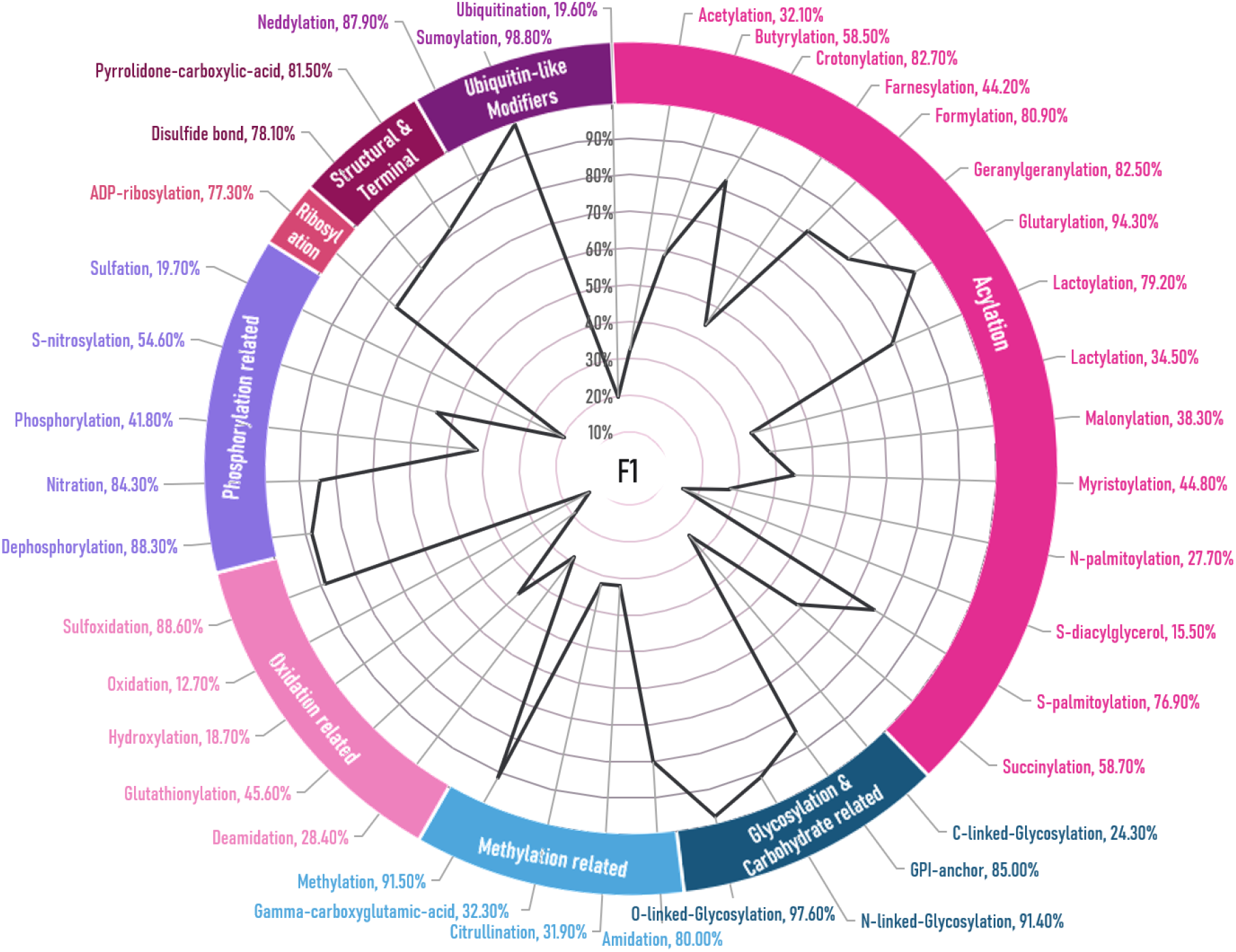
Macro-F1 values for 39 PTM labels predicted by AstraPTM2

- **High performance (***>***80% F1):** Many rare or motif-driven PTMs such as methylation (F1=91.5%), O-linked glycosylation (97.6%), sumoylation (98.8%) and neddylation (87.9%).
- **Moderate (30–60% F1):** Succinylation (58.7%), butyrylation (58.5%), S-nitrosylation (54.6%), glutathionylation (45.6%).
- **Low-to-moderate (***<***40% F1):** Broad, heterogeneous PTMs such as phosphorylation (41.8%), acetylation (32.1%), and ubiquitination (19.6%).

Some of the lowest scores are seen for common PTMs such as phosphorylation, which are the most widely evaluated in the literature. This reflects AstraPTM2’s goal of maximally broad coverage of multiple PTM types, rather than being tuned to a single common PTM family. Along with that, not specifically focusing on a certain family of proteins or certain type of PTMs.

**Table 2:**
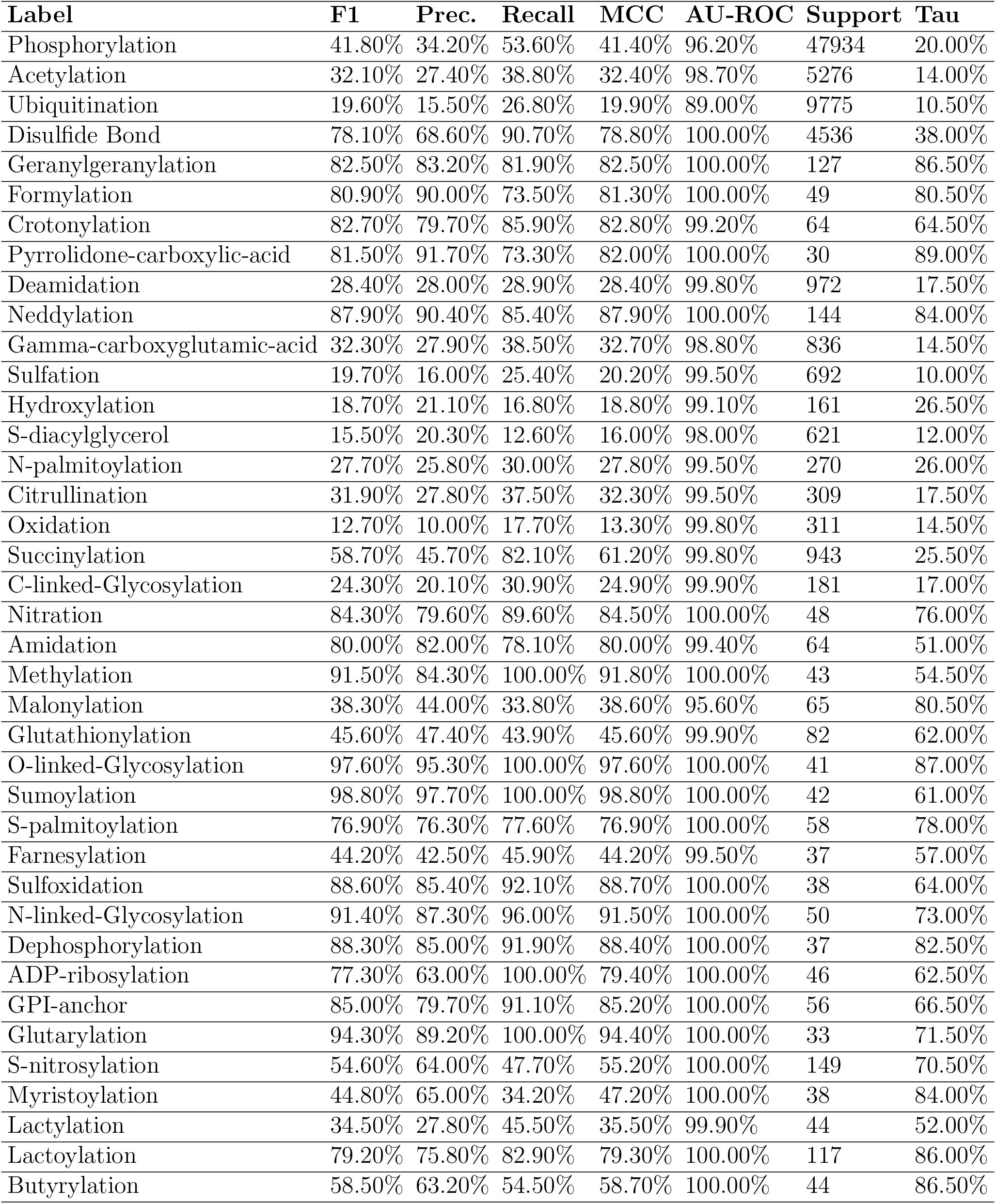
Hold-out test evaluation results for 39 PTM labels.

| **Label** | **F1** | **Prec.** | **Recall** | **MCC** | **AU-ROC** | **Support** | **Tau** |
| --- | --- | --- | --- | --- | --- | --- | --- |
| Phosphorylation | 41.80% | 34.20% | 53.60% | 41.40% | 96.20% | 47934 | 20.00% |
| Acetylation | 32.10% | 27.40% | 38.80% | 32.40% | 98.70% | 5276 | 14.00% |
| Ubiquitination | 19.60% | 15.50% | 26.80% | 19.90% | 89.00% | 9775 | 10.50% |
| Disulfide Bond | 78.10% | 68.60% | 90.70% | 78.80% | 100.00% | 4536 | 38.00% |
| Geranylgeranylation | 82.50% | 83.20% | 81.90% | 82.50% | 100.00% | 127 | 86.50% |
| Formylation | 80.90% | 90.00% | 73.50% | 81.30% | 100.00% | 49 | 80.50% |
| Crotonylation | 82.70% | 79.70% | 85.90% | 82.80% | 99.20% | 64 | 64.50% |
| Pyrrolidone-carboxylic-acid | 81.50% | 91.70% | 73.30% | 82.00% | 100.00% | 30 | 89.00% |
| Deamidation | 28.40% | 28.00% | 28.90% | 28.40% | 99.80% | 972 | 17.50% |
| Neddylation | 87.90% | 90.40% | 85.40% | 87.90% | 100.00% | 144 | 84.00% |
| Gamma-carboxyglutamic-acid | 32.30% | 27.90% | 38.50% | 32.70% | 98.80% | 836 | 14.50% |
| Sulfation | 19.70% | 16.00% | 25.40% | 20.20% | 99.50% | 692 | 10.00% |
| Hydroxylation | 18.70% | 21.10% | 16.80% | 18.80% | 99.10% | 161 | 26.50% |
| S-diacylglycerol | 15.50% | 20.30% | 12.60% | 16.00% | 98.00% | 621 | 12.00% |
| N-palmitoylation | 27.70% | 25.80% | 30.00% | 27.80% | 99.50% | 270 | 26.00% |
| Citrullination | 31.90% | 27.80% | 37.50% | 32.30% | 99.50% | 309 | 17.50% |
| Oxidation | 12.70% | 10.00% | 17.70% | 13.30% | 99.80% | 311 | 14.50% |
| Succinylation | 58.70% | 45.70% | 82.10% | 61.20% | 99.80% | 943 | 25.50% |
| C-linked-Glycosylation | 24.30% | 20.10% | 30.90% | 24.90% | 99.90% | 181 | 17.00% |
| Nitration | 84.30% | 79.60% | 89.60% | 84.50% | 100.00% | 48 | 76.00% |
| Amidation | 80.00% | 82.00% | 78.10% | 80.00% | 99.40% | 64 | 51.00% |
| Methylation | 91.50% | 84.30% | 100.00% | 91.80% | 100.00% | 43 | 54.50% |
| Malonylation | 38.30% | 44.00% | 33.80% | 38.60% | 95.60% | 65 | 80.50% |
| Glutathionylation | 45.60% | 47.40% | 43.90% | 45.60% | 99.90% | 82 | 62.00% |
| O-linked-Glycosylation | 97.60% | 95.30% | 100.00% | 97.60% | 100.00% | 41 | 87.00% |
| Sumoylation | 98.80% | 97.70% | 100.00% | 98.80% | 100.00% | 42 | 61.00% |
| S-palmitoylation | 76.90% | 76.30% | 77.60% | 76.90% | 100.00% | 58 | 78.00% |
| Farnesylation | 44.20% | 42.50% | 45.90% | 44.20% | 99.50% | 37 | 57.00% |
| Sulfoxidation | 88.60% | 85.40% | 92.10% | 88.70% | 100.00% | 38 | 64.00% |
| N-linked-Glycosylation | 91.40% | 87.30% | 96.00% | 91.50% | 100.00% | 50 | 73.00% |
| Dephosphorylation | 88.30% | 85.00% | 91.90% | 88.40% | 100.00% | 37 | 82.50% |
| ADP-ribosylation | 77.30% | 63.00% | 100.00% | 79.40% | 100.00% | 46 | 62.50% |
| GPI-anchor | 85.00% | 79.70% | 91.10% | 85.20% | 100.00% | 56 | 66.50% |
| Glutarylation | 94.30% | 89.20% | 100.00% | 94.40% | 100.00% | 33 | 71.50% |
| S-nitrosylation | 54.60% | 64.00% | 47.70% | 55.20% | 100.00% | 149 | 70.50% |
| Myristoylation | 44.80% | 65.00% | 34.20% | 47.20% | 100.00% | 38 | 84.00% |
| Lactylation | 34.50% | 27.80% | 45.50% | 35.50% | 99.90% | 44 | 52.00% |
| Lactoylation | 79.20% | 75.80% | 82.90% | 79.30% | 100.00% | 117 | 86.00% |
| Butyrylation | 58.50% | 63.20% | 54.50% | 58.70% | 100.00% | 44 | 86.50% |

### Comparator Models

Against its immediate predecessor AstraPTM, AstraPTM2 improves F1 by 9–14 pp and expands coverage from 25 to 39 PTMs. Multiple other PTM prediction methods have been published in the past few years, utilising a variety of architectures (see Table 3).

**Table 3:** Key properties of multi-PTM prediction models published 2023–2025.

| Model (Year) | # Labels | Dataset Size / Seq. Limit | Architecture | Notes |
| --- | --- | --- | --- | --- |
| **AstraPTM2 (2025)** | **39** | >100K proteins, no length limit | Transformer + AF2 + convs | Context-aware, calibrated |
| AstraPTM (2025) | 25 | 75K proteins, no length limit | Transformer, dual heads | First-gen model |
| UniPTMs (16) (2025) | 5 | Not reported, 1022 aa | Dual-path multimodal | Focused on high-confidence PTMs |
| PDeepPP (2025) | ~13 | 33-aa sliding window | Hybrid transformer+CNN | Peptide-based prediction |
| PTM-Mamba (2025) | 25 | ~80K proteins, 1200 aa | PLM + Mamba blocks | Focused on phospho/ubiquitin |
| MTPrompt-PTM (2025) | 13 | UniProt, 1022 aa | Prompt-tuned PLM | Truncated proteins |
| DeepMVP (2025) | 6 | 397K PTM sites, 31–61 aa windows | Ensemble models | MS reprocessed dataset |
| MeToken (2024) | 25 | Not reported | Seq+struct tokenization | Limited benchmark coverage |
| PTM-CMGMS (2024) | n/a | Not reported | Contrastive learning | Tested on a few PTMs only |
| PTM-GPT2 (2024) | 19 | Not reported, 1022 aa | GPT-2 decoder | ProtGPT2 fine-tuning |
| SiteTACK (2024) | 13 | Not reported | CNN + PTM encoding | Focused on 13 PTMs |
| MIND-S (2023) | 13 | Not reported | Multi-label seq+struct | Multi-task design |

With macro-F1=59% and AUROC=0.99, AstraPTM2 shows strong performance against other multi-label PTM predictors, however direct comparisons are hindered by heterogenous label use and different metrics reported. Metrics and benchmarks used for model assessment also vary widely, with some model publications lacking clarity on specifics.Despite this, in comparison of closest available metrics (e.g. for a subset of AstraPTM2 labels) AstraPTM2 showed comparable or stronger performance than the comparator models. Notably, AstraPTM2 performed somewhat worse on the most common PTM labels that were most frequently evaluated by other models, indicating an even bigger advantage for a diverse set of less common PTM labels.

In addition, many models only evaluated small windows (e.g. 33 residue peptides in PDeepPP (6) and 31–61 residues in DeepMVP (3)) or substantially limited overall protein length in the database. In contrast, AstraPTM2 employed a self-attention architecture without any window limitations.

The number of PTM labels in AstraPTM2 is particularly noteworthy, especially since AstraPTM2 labels were typically less granular than those of comparator models: while other models frequently separated one PTM into several labels based on the amino acid targeted, AstraPTM2 only explicitly separated glycosylation and palmitoylation labels. This underscores that the high number of labels covered by AstraPTM2 is really reflective of a large variety of included PTM types and is not artificially increased by granularity.

**Table 4:** Performance comparison of AstraPTM2 with other multi-PTM predictors.

| Model (Year) | Reported Metric(s) | Closest AstraPTM2 Metric | Comparison Outcome |
| --- | --- | --- | --- |
| **AstraPTM2 (2025)** | Macro-F1=59%, AUROC=0.99 | — | — |
| AstraPTM (2025) | Macro-F1=49% | Macro-F1=59% | **+10 pp**, +14 PTMs |
| UniPTMs (2025) | Micro-MCC=44–79% | Micro-MCC=41–98% | Comparable MCC, much wider coverage |
| PDeepPP (2025) | Micro-MCC=59–98% | Micro-MCC=20–99% | Similar MCC, AstraPTM2 full-length seq |
| PTM-Mamba (2025) | Phospho F1 <20% | Phospho F1=42% | **2× better phospho F1** |
| MTPrompt-PTM (2025) | F1=77–93% (subset) | F1=20–99% (subset) | Higher coverage, similar quality on overlap |
| DeepMVP (2025) | AUROC >0.85 | AUROC >0.89 | Comparable AUROC, no window limit |
| MeToken (2024) | F1 ~50% | F1=59% | +9 pp improvement |
| PTM-CMGMS (2024) | MCC=31–70% | MCC=38–83% | Higher MCC on overlap PTMs |
| PTM-GPT2 (2024) | Micro-MCC=16–90% | Micro-MCC=20–99% | Similar range, broader PTM set |
| SiteTACK (2024) | AUROC=0.90–0.99 | AUROC=0.89–1.00 | Comparable AUROC, +26 PTMs |
| MIND-S (2023) | Macro-MCC=73% | Macro-F1=59% | Comparable on overlap, 3× more PTMs |

## User Experience

### End-to-end flow

AstraPTM2 is integrated into the Orbion web platform. Users input a protein sequence, which triggers the computational pipeline: embeddings and AlphaFold2 structures are generated, followed by residue-level PTM scoring in both *calibrated* and *global-threshold* modes. Results are displayed in synchronized 2D and 3D views.

### 2D & 3D visualization

The 2D view presents per-residue blocks with PTM types listed on the left, including badges for high- and low-confidence sites. The 3D viewer (implemented using the <monospace>3Dmol.js</monospace> library)(14) overlays annotations directly on the protein structure, with tooltips showing label information. These complementary views allow users to quickly spot patterns along the sequence and reason about their spatial distribution within the folded protein.

**Figure 5:**
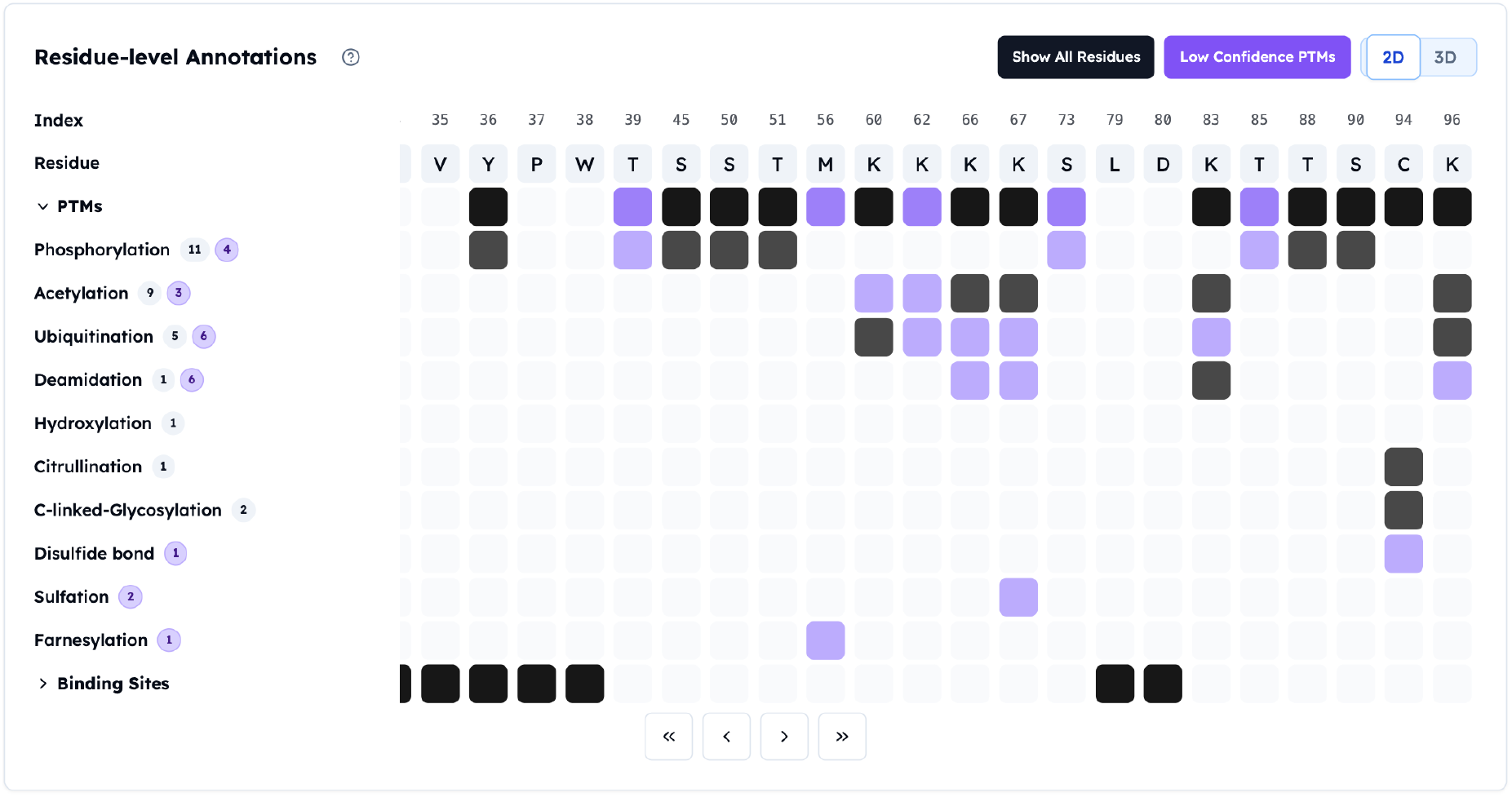
2D view of AstraPTM2 predictions for Hemoglobin Subunit Beta in the Orbion web platform. High-confidence sites are shown in black; low-confidence sites are shown in violet.

**Figure 6:**
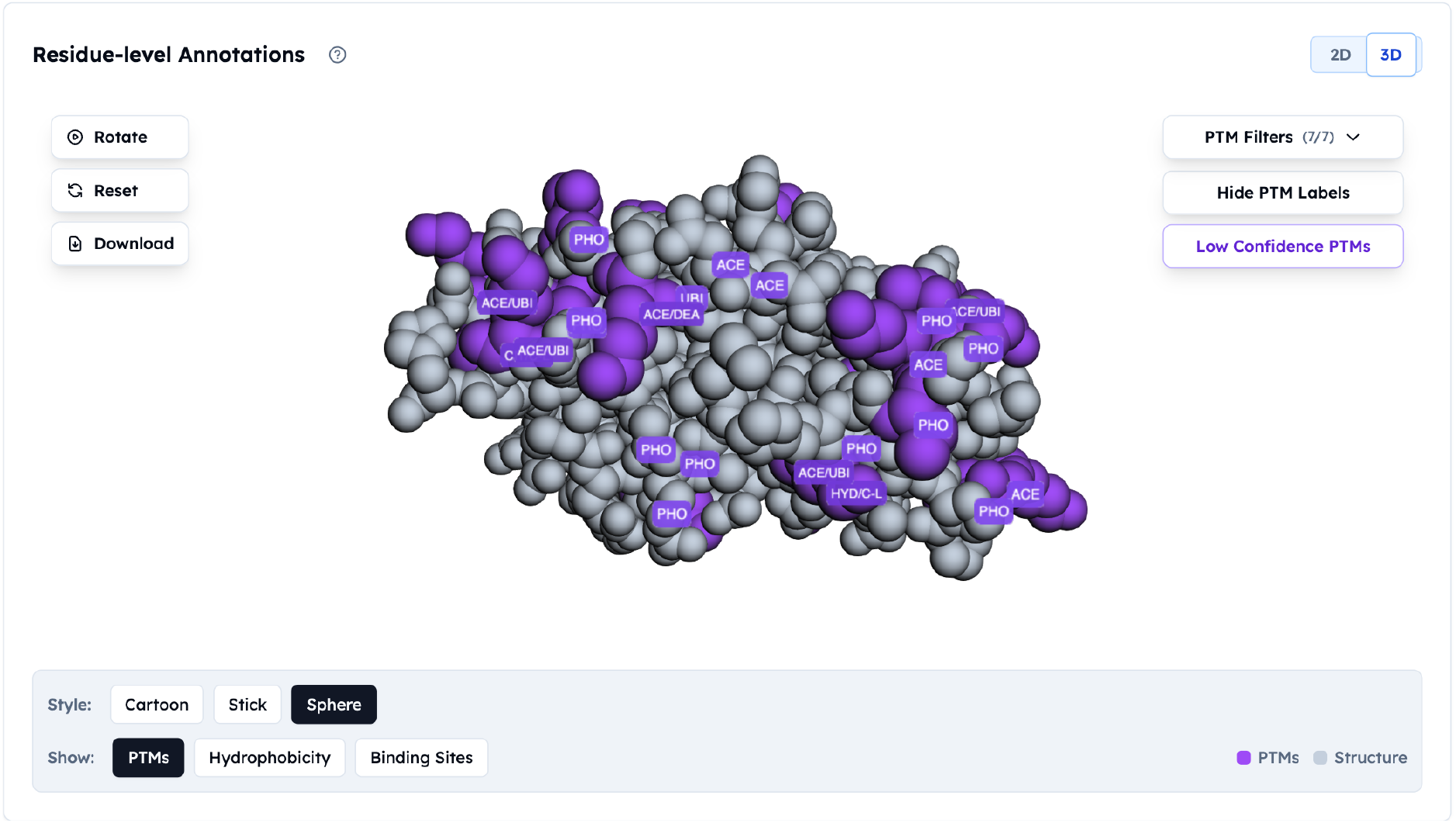
3D view of AstraPTM2 predictions for Hemoglobin Subunit Beta using 3Dmol.js. Only high-confidence sites are displayed for clarity.

### Dual operating modes

Two complementary modes are provided:

#### 1. Calibrated mode

Uses per-label affine calibration and held-out thresholds (*τ*_ℓ_) for actionable, high-specificity predictions.

#### 2. Exploration mode

Applies a single global threshold to surface lower-confidence or under-documented sites, enabling hypothesis generation in poorly annotated regions.

Both modes are displayed side-by-side and encoded in downloads for reproducibility.

### Export and provenance

One-click export generates a .pdb file with PTM annotations in <monospace>B-factor</monospace> or <monospace>REMARK</monospace> fields. Each export embeds model version, calibration set ID, and timestamp to ensure reproducibility.

### Interpretation guidelines

Use calibrated mode for downstream decision-making (e.g., mutagenesis planning). Use exploration mode for hypothesis generation and prioritizing follow-up experiments in poorly characterized regions.

## Example Predictions

To illustrate AstraPTM2 on well-studied, mechanistically diverse proteins, we evaluated five canonical targets. Across GPCR, hormone, tumor suppressor, phosphatase, and chaperone classes, AstraPTM2 recovers hallmark PTMs (e.g., GPCR C-tail phosphorylation clusters, insulin disulfide bonds, PTEN C-terminal phosphorylation) while surfacing additional plausible sites that may merit follow-up. High-confidence sites are determined by per-label calibrated thresholds.

**Table 5:** Representative AstraPTM2 predictions (UniProt canonical indices).

| Protein (UniProt) | Predicted PTMs (count & residue indices) | Notes / High-confidence sites* |
| --- | --- | --- |
| **Rhodopsin (P08100)** | *Phosphorylation* (6): 333, 335, 337, 339, 341, 342<br>*Ubiquitination* (1): 338<br>*Disulfide bond* (2): 109, 186<br>*Citrullination* (2): 321, 322<br>*Succinylation* (1): 14<br>*ADP-ribosylation* (1): 337 | High-confidence: several C-tail phosphorylation sites (>0.4), disulfide at 109; classic GPCR regulation motif recapitulated. |
| **Insulin (P01308)** | *Disulfide bond* (6): 30, 42, 94, 95, 99, 108 | All six disulfides >0.65, matching A-B chain connectivity; strong structural sanity check. |
| **Cellular Tumor Antigen p53 (P04637)** | *Phosphorylation* (13): 93, 302, 311, 312, 313, 314, 326, 361, 365, 366, 375, 386, 391<br>*Acetylation* (1): 385<br>*Ubiquitination* (2): 119, 350<br>*Hydroxylation* (1): 123<br>*N-palmitoylation* (1): 385 | High-confidence: multiple Ser/Thr sites (>0.45) in transactivation/CTD regions; lysine 385 flagged for acetylation. |
| **PTEN (P60484)** | *Phosphorylation* (20): 166, 228, 293, 337, 354, 359, 360, 361, 362, 363, 365, 369, 376, 378, 379, 381, 382, 384, 397, 400<br>*Ubiquitination* (11): 59, 79, 101, 146, 162, 182, 236, 288, 312, 321, 401 | High-confidence: nine phosphorylation sites (>0.3) in the C-terminal tail; multiple lysines consistent with reported stability control. |
| **Heat shock protein HSP 90-alpha (P07900)** | *Phosphorylation* (4): 67, 230, 251, 262<br>*Acetylation* (34): 40, 57, 68, 73, 184, 190, 227, 254, 282, 291, 313, 326, 361, 406, 409, 430, 434, 435, 442, 477, 484, 488, 498, 533, 538, 545, 566, 572, 575, 580, 584, 653, 656, 659<br>*Ubiquitination* (25): 68, 111, 184, 208, 227, 254, 282, 291, 293, 313, 326, 361, 406, 430, 435, 442, 445, 477, 488, 533, 538, 545, 572, 575, 614<br>*Hydroxylation* (3): 373, 528, 596 | High-confidence: acetylation and ubiquitination clusters across middle/CTD regions; phosphorylation at 230 and 262 near functional sites. |

\*High-confidence refers to per-label calibrated thresholds derived on held-out data. Predictions are *in silico* and intended to guide experimental design.

## Discussion

AstraPTM2 achieves state-of-the-art macro-F1 (59%) while covering 39 distinct PTMs, making it one of the most comprehensive residue-level PTM predictors to date. Its development highlighted several key principles that go beyond raw model capacity and may serve as design guidelines for future multi-PTM predictors.

Reliable prediction of PTM sites and types across a broad spectrum remains challenging. Experimental PTM data are noisy, incomplete, and highly context-dependent, with many “negatives” simply representing untested sites. While some models focus narrowly on the most common PTMs using curated resources such as UniProt (15) or PTMAtlas (3), AstraPTM2 was built to generalize across rare and common PTMs, enabling both discovery and hypothesis generation.

Several general insights emerged from the AstraPTM2 development process:

- **Calibration is critical**. Per-label affine calibration and *β*-smoothed threshold search were decisive in achieving stable, fair F1 scores across PTM groups. Without calibration, results fluctuated across runs and biased toward abundant PTMs.
- **Curriculum learning is a major lever**. Staging the training (common → medium rare→ labels) with a balanced schedule (e.g., 25–15–15 epochs) consistently improved generalization. Overextending the rare-label stage led to overfitting, while under-emphasizing it caused collapse in rare-label recall.
- **Motif-level modeling is essential**. Adding local context through depth-wise convolutions (3×1, 5×1, 7×2) and motif-mixing branches produced the single largest macro-F1 improvements (+2–3 pp each). These layers captured short motifs like kinase target sites that attention alone under-represented.
- **Balanced data beats more data**. Merely scaling to a 230k-protein dataset degraded performance. Stratified, balanced subsets of ~ 100k proteins with rare-label oversampling consistently outperformed unbalanced larger corpora.
- **Auxiliary supervision enhances rare-label learning**. Kinase-group classification and site-window auxiliary tasks improved representation learning and gave the highest lift for labels with *<*100 sites in training data.
- **Negative sampling must be tuned**. Both oversampling negatives and undersampling them harmed model stability. A dynamic NEG SAMPLING RATIO in the 1.0–1.5 range yielded optimal results.
- **Architectural experiments clarified priorities**. Multi-head groupings (common, medium, rare) and learnable temperature multipliers improved balance between frequent and rare PTMs, whereas overly complex modules (e.g., gated MoE heads, asymmetric focal loss) often hurt convergence. Lightweight structural branches (GCN/GAT) added modest gains (~ +0.3–0.5 pp) but were secondary to sequence-local modeling.
- **Hyperparameter tuning had non-linear effects**. Sweeping focal-loss *γ* per-label, adjusting learning-rate schedules, and applying short warm-restart cycles improved training stability. However, more exotic approaches (e.g., PU loss, class-balanced focal) underperformed compared to adaptive focal loss with decayed *γ*.

Together, these findings indicate that balanced curation of data, thoughtful curriculum scheduling, and motif-sensitive architectures are far more impactful than simply scaling model size or dataset volume. Importantly, calibrated thresholds and dual prediction modes make AstraPTM2’s outputs easier to interpret biologically, enabling researchers to prioritize high-confidence sites for mutagenesis while still exploring low-confidence predictions for hypothesis generation.

## Conclusions

AstraPTM2 represents a step-change in residue-level PTM prediction: it is, to our knowledge, the first model to provide calibrated, context-aware predictions for 39 distinct PTM types across full-length protein sequences. By combining ESM-2 embeddings, AlphaFold2-derived structural features, and motif-sensitive convolutions within a transformer backbone, AstraPTM2 achieves macro-F1 of 59% and AUROC of 0.99—competitive with or stronger than recently published multi-PTM predictors despite its much broader label coverage.

Beyond performance metrics, AstraPTM2 demonstrates that:

- Broad PTM coverage can be achieved without sacrificing predictive power, provided that training data are balanced and curriculum learning is carefully staged.
- Motif-level modeling and per-label calibration substantially improve interpretability, enabling users to make actionable decisions (e.g., site-directed mutagenesis, expression construct design) with confidence.
- Dual-mode outputs (calibrated vs. exploratory) empower researchers to distinguish high-confidence actionable sites from lower-confidence hypothesis-generating predictions, facilitating wet-lab validation strategies.

AstraPTM2 sets a new benchmark for broad-spectrum PTM prediction, combining high coverage, calibrated interpretability, and deployment-ready outputs. We believe its design - balanced curation, curriculum learning, motif-aware encoders - can guide the next generation of multi-PTM predictors and accelerate experimental discovery.

